# Targeting Ligand Shape While Preserving a Warhead through Guided Diffusion and Adaptive Staged Growth

**DOI:** 10.64898/2026.07.21.739744

**Authors:** Ismael Castañón

## Abstract

Fragment-based growth models exploit pre-existing structures but can become constrained by early autoregressive decisions and generally lack knowledge of the final target shape. Here, we tested whether geometric guidance applied to a pretrained DiffSBDD model could steer generation toward the shape of a ligand B while preserving a warhead from ligand A. One-shot guidance increased target-volume coverage but predominantly produced disconnected structures. Reformulating the task as smaller scaffold expansions substantially improved the recovery of connected anchor components. Target-directed selection and adaptive growth increments enabled progressive movement toward B, while beam search improved endpoint recovery. Soft-scaffold inpainting provided graded control over the mobility of the inherited scaffold while preserving the fixed warhead. Together, these results show that inference-time shape guidance can be converted from disconnected volume coverage into controlled staged molecular growth, although performance remains dependent on the geometric compatibility of the ligand pair.

## INTRODUCTION

Three-dimensional generative models based on equivariant neural networks make it possible to design ligands conditioned on a known pocket, generating structures that are plausible given its geometry. Diffusion and flow-matching models have gained prominence in recent years thanks to their capacity to model three-dimensional molecular distributions through iterative denoising ^1–3^. This structural flexibility, however, comes at the price of generating molecules that are less synthesizable and that adopt improbable atomic arrangements, showing an absence of rings, poorly coordinated atoms, and unfavorable geometric expansions^4^. Computational chemistry, on the other hand, commonly requires strategies that exploit knowledge from prior ligands, such as preserving known substructures and building stable scaffolds. For this reason, numerous frameworks have emerged that focus on local inpainting or on the connectivity of different structures ^5^. The best-known approaches are the autoregressive growth of atoms or fragments using GNNs, although growth via MCTS has also been proposed ^6–8^. Such approaches may nonetheless be suboptimal for tasks requiring global shape remodeling, because a fixed set of building blocks imposes a discrete granularity that can bias the search or trap growth in locally favorable trajectories. More generally, autoregressive and diffusion-based generators exhibit complementary trade-offs: sequential methods provide greater control over connectivity and local chemical structure, whereas diffusion models can account for the global three-dimensional context more directly. A different strategy is the one adopted by AutoFragDiff ^9^, a diffusion model trained on retrosynthetic routes that performs autoregressive growth through a diffusion model conditioned on fragment-based elaboration. This approach combines both paradigms, giving it the flexibility of diffusion models and the robustness of fragment-based methods. It nevertheless still presents the problems associated with autoregressive growth, and, like other purpose-trained models, it requires a dedicated training procedure rather than reusing a general-purpose generator.

An alternative to retraining is to steer a pretrained model at inference time. Rather than learning a new conditional distribution, inference-time guidance perturbs the sampling trajectory of an existing model toward a desired property, for example by adding a gradient derived from an external objective at each denoising step. This makes it possible to impose constraints that the model was never explicitly trained on, at the cost of pushing the sampler away from the region of the prior where its outputs are most reliable. In structure-based design, one such objective is molecular shape: the three-dimensional volume a ligand occupies governs both its complementarity to the pocket and its three-dimensional similarity to known binders, which makes shape overlap a natural target for guidance when a reference molecule is available.

In this work we set out to test whether a general-purpose diffusion model would be able to preserve a local motif A while generating the shape of a molecule B, using this local motif as a hard constraint and the shape of B as a soft, inference-time guidance signal. The rationale behind this hypothesis is a phenomenon commonly observed in generative frameworks: although certain models have a remarkable capacity to generalize and classify, they perform poorly when they must generate from the prior without a seed. Anchoring the generation on a known motif provides such a seed, while the shape of a second molecule supplies a direction in which to grow. This would also make it possible to combine pre-existing knowledge from two different candidates, a situation that arises frequently in the pharmaceutical industry, where fragments or scaffolds from separate hits are merged, in order to generate a new molecule that better exploits both structures.

The main problem with this approach is that geometric guidance imposed on the prior tends to generate disconnected molecules. Over the course of this project, however, we have shown how a fragment-based growth strategy can be used to steer generation using shape-based metrics while keeping the growing molecule connected. This approach, on the other hand, depends heavily on the compatibility between the geometries of the two structures, which opens a new problem of selection and optimization.

## METHODS

### Structural systems and dual-conditioning task

Reference ligands were obtained from the crystallographic SARS-CoV-2 Mpro fragment-screening dataset generated through the Diamond XChem platform and subsequently used in the SILVR study ^11^. All structures were prepared in a common PanDDA-aligned receptor coordinate frame, allowing direct comparison of ligand coordinates without post hoc molecular alignment ^12^.

Two ligand pairs were studied. The x0434-to-x2193 pair is referred to as the hard pair, whereas x0874-to-x1093 is referred to as the moderate pair. In each pair, ligand A supplied the local substructure to be preserved and ligand B supplied the global three-dimensional shape target. A seven-heavy-atom substructure of ligand A was used as the fixed warhead. The warhead was defined independently of ligand B and was not adapted to the target shape.

The geometric compatibility of each pair was measured directly in the common receptor frame using RDKit Shape Tanimoto distance and directional Shape Protrude distance. Both quantities are distances, such that lower values indicate greater geometric similarity. The moderate pair had an A-to-B Shape Tanimoto distance of 0.728 and a Shape Protrude distance of 0.500. The hard pair had an A-to-B Shape Tanimoto distance of 0.874 and a Shape Protrude distance of 0.769.

### Frozen diffusion model and inference-time shape-guidance

#### Pocket-conditioned diffusion and hard warhead inpainting

Molecules were generated using a frozen pocket-conditioned DiffSBDD checkpoint operating on ligand atom types and Cartesian coordinates in the receptor frame. No model parameters were updated during the study.

Let *z*_*t*_ denote the noisy ligand state at reverse-diffusion step *t*, including coordinate and atom-feature channels, and let *P* denote the protein-pocket representation. The frozen denoiser predicts a clean ligand state according to

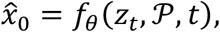

where *θ* denotes the fixed model parameters.

The local condition was imposed through molecular inpainting. A binary mask *M*_*fix*_ identified the atoms belonging to the current fixed scaffold, and

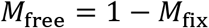

identified the atoms generated at the current stage. During every reverse-diffusion step, the fixed atoms were sampled from the appropriately noised representation of the supplied scaffold and reintroduced through the inpainting mask. The generated coordinates therefore could not permanently overwrite the local scaffold.

In the initial stage, the fixed scaffold consisted of the seven-heavy-atom warhead from A. In subsequent staged-growth steps, the complete selected scaffold from the preceding stage was supplied as the fixed structure.

#### Differentiable Gaussian shape objective

Global guidance was applied using a differentiable Gaussian approximation to volumetric Shape Tanimoto similarity. For two sets of heavy-atom coordinates *X* and *Y*, their Gaussian overlap was defined as

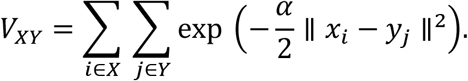

Let *G* denote the complete predicted ligand, including both fixed and free atoms, and let *B* denote the heavy-atom coordinates of the target ligand in the same receptor frame. The differentiable shape score was

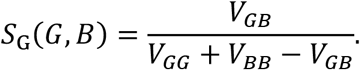

The cross-overlap term *V*_*GB*_ rewards occupation of the volume defined by B. The self-overlap term *V*_*GG*_ introduces an opposing contribution that discourages all generated atoms from collapsing into the same spatial region.

The fixed atoms contributed to the ligand volume in *V*_*GG*_ and *V*_*GB*_, but gradients were calculated only with respect to the free coordinates:

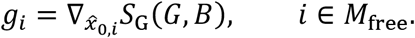

The Gaussian width was set to

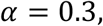

which had previously been calibrated to produce a smooth overlap scale consistent with the independent RDKit grid-based evaluator.

#### Guidance integration into the reverse process

The gradient was clipped independently for each free atom while preserving its direction:

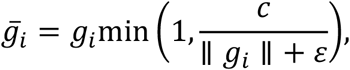

where *c* = 1.0 Å was the per-atom gradient-norm threshold.

To deform the predicted molecular shape without translating the complete set of free atoms, the mean guidance vector was removed:

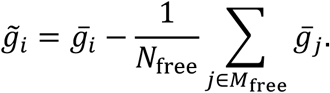

The coordinate displacement requested by the global guidance was then

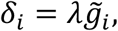

where λ is the experimental guidance strength.

Guidance was applied in predicted-clean-coordinate space rather than directly to the noisy state. The displacement of 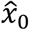 was introduced into the diffusion sampler through the corresponding correction to the predicted noise:

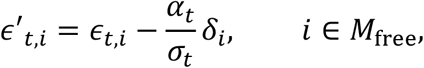

where α_*t*_ and σ_*t*_ are the diffusion signal and noise coefficients at step *t*. The reverse-transition mean was subsequently calculated from ϵ′_*t*_. Atom-feature channels were not modified by the shape gradient.

The sequence of operations at each guided reverse step was therefore: denoiser prediction, shape-gradient evaluation on the free coordinates, gradient clipping and center-of-mass neutralization, correction of the predicted noise, and hard inpainting of the fixed scaffold.

The hard scaffold overwrite was consequently the final conditioning operation. The global objective could bias the generated atoms but could not displace the fixed scaffold.

Unless otherwise stated, sampling used 50 diffusion timesteps and five resampling iterations.

### Staged fragment-growth framework

#### Recursive scaffold expansion

Staged growth reformulated the dual-conditioning task as a sequence of smaller molecular expansions. Every trajectory began from the bare seven-heavy-atom warhead.

At stage *s*, the current scaffold *C*_*s*_ was supplied to DiffSBDD through the inpainting interface. All atoms of *C*_*s*_ were treated as fixed, and a requested number of additional atoms, denoted by *n*_add_, was generated around it under shape guidance toward B. The model produced a set of candidate records

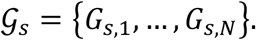

Each record was chemically sanitized, decomposed into connected components, and evaluated. One or more valid components were selected and propagated as the fixed scaffolds of stage *s* + 1.

The global target B remained unchanged throughout the trajectory. Staging therefore modified the size of the requested generative action but not the direction of the shape guidance.

#### Warhead identification and anchor component

The seven warhead atoms were identified in each generated record by element-sensitive geometric matching to their coordinates in ligand A. A warhead match was accepted when every fixed atom lay within 0.2 Å of its reference position.

The anchor component was defined as the connected component containing all seven matched warhead atoms. A record was assigned no valid anchor when one or more warhead atoms could not be matched, when the maximum displacement exceeded 0.2 Å, or when the matched atoms occurred in different connected components.

The heavy-atom count of the anchor component was denoted by

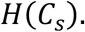

A generated record was considered fully connected only when all its heavy atoms belonged to a single connected component. Full-record connectivity and anchor-component validity were recorded separately. A fragmented record could therefore contain a valid growing anchor together with one or more secondary disconnected fragments.

Only the anchor component was eligible for propagation. Secondary fragments were never incorporated into the scaffold of the following stage.

#### Component-aware shape evaluation

Shape metrics were calculated on the anchor component rather than on the complete generated record. This prevented disconnected secondary components from artificially improving volumetric overlap with B.

RDKit Shape Tanimoto distance was calculated against both A and B:

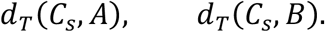

Shape Protrude distance was similarly calculated as

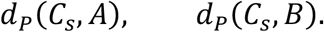

Hydrogens were ignored. All metrics were calculated directly in the common receptor coordinate frame without molecular alignment. Shape Protrude was evaluated by preserving the direction from the generated anchor component to the reference ligand when the compared structures differed in size.

Shape Tanimoto distance is symmetric and measures overall volumetric disagreement. Shape Protrude distance is asymmetric and measures the fraction of the generated anchor volume extending beyond the selected reference volume. Lower values indicate a closer match in both cases.

For a trajectory beginning at stage *s*_0_ and terminating at stage *s*_*f*_, Shape Tanimoto improvement toward B was defined as

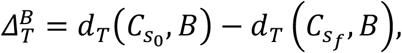

and Shape Protrude improvement as

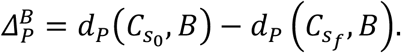

Positive values indicate movement toward B.

### Experimental design

#### Single-shot reference campaign

The complete atom budget was generated in one inpainting call around the seven-atom warhead. Both ligand pairs were evaluated at guidance strengths.

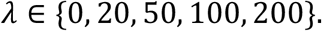

Five random seeds were used for each pair and guidance strength, and each run requested ten samples.

The campaign measured the connected-molecule rate, number of heavy-atom-containing components, heavy-atom fraction retained in the anchor component, and the difference between full-record and component-aware shape scores.

#### Parameterization of staged growth

A factorial parameter sweep was performed to determine how the local size of each generative action affected connectivity and shape. The per-stage increment was varied over:

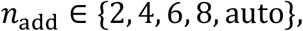

Where auto mode allowed DiffSBDD to select the number of added atoms.

The number of resampling iterations was varied over:

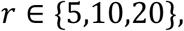

and the shape-guidance strength was varied over:

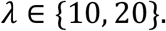

The complete grid was evaluated for both ligand pairs. Each record was audited for sanitization, warhead retention, full-record connectivity, anchor-component size, and anchor-component shape relative to A and B.

This sweep was descriptive. All conditions used shape guidance toward B, and no B-blind selection contrast was included. It was therefore used to characterize the effects of increment size, resampling, and guidance strength, rather than to attribute movement toward B to directed scaffold selection.

#### Controlled fixed-increment staged growth

A controlled multi-stage campaign was performed to determine whether explicit selection relative to ligand B produced directional improvement beyond the effects of staged sampling and candidate multiplicity.

Four fixed per-stage increments were evaluated:

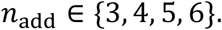

The +3 and +4 arms contained three growth stages, whereas the +5 and +6 arms contained two stages, producing a total requested growth of 9-12 atoms beyond the initial seven-heavy-atom warhead.

Three propagation regimes were compared. The single-sample control, denoted A1, generated one candidate at each stage. When the candidate contained a valid anchor component, that component was propagated without consulting ligand B. Because no alternative candidate was available, A1 measured the behavior and robustness of unselected sequential growth.

The best-of-ten B-blind control, denoted A10, generated ten candidates at each stage. Among candidates containing a valid anchor, the anchor component with the greatest heavy-atom count was propagated. Ligand B was not used in the selection rule. A10 therefore measured the benefit of candidate multiplicity and structural selection without target-directed selection.

The directed branch also generated ten candidates at each stage. Candidate selection consulted ligand B. A candidate was eligible for propagation when neither its Shape Tanimoto distance nor its Shape Protrude distance to B increased relative to the parent scaffold. Among eligible candidates, the candidate producing the greatest joint reduction in the two distances was propagated.

The primary controlled comparison was between A10 and the directed branch. Both generated ten candidates per stage and differed only in whether B was used during scaffold selection, thereby isolating the contribution of target-directed selection from the best-of-ten sampling advantage. A1 served as an auxiliary control for the robustness of single-sample staged growth.

Ten base seeds were evaluated for each ligand pair and increment arm. A1 and A10 each used three independently derived replicate seeds per base seed, yielding 240 trajectories per branch. The directed branch used one trajectory per base seed, yielding 80 trajectories. The complete campaign therefore comprised 560 trajectories.

The guidance strength was fixed at λ = 20, with five resampling iterations and 50 diffusion timesteps.

A trajectory completed when it reached the prespecified number of stages for its increment arm. It terminated earlier when a stage failed to produce a valid anchor, when molecular generation did not complete successfully, or, in the directed branch, when none of the generated candidates satisfied the joint shape-progression criterion. Early termination was retained as a trajectory-level outcome rather than treated as missing data.

Because the A1 and A10 branches used independently derived replicate seeds whereas the directed branch used the base seeds, comparisons between A10 and directed selection were performed at the ligand-pair-by-increment level rather than as paired trajectory comparisons.

### Adaptive search over staged growth

The final growth campaign treated the per-stage atom increment as a decision variable rather than as a fixed schedule.

At every stage, each live search state generated candidates under all increments:

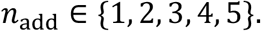

Ten samples were drawn for every increment. Candidates generated from all live parents were audited jointly.

Two variants were evaluated: greedy search, with beam width *k* = 1, and beam search, with beam width *k* = 3. The two variants shared the same candidate-generation procedure, validity criteria, progression gate, target size, and ranking function. Beam width was therefore the only intended algorithmic difference.

For a parent scaffold *C*_*p*_ and candidate child *C*_*c*_, local gains were defined as

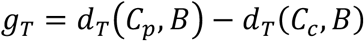

and

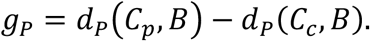

A candidate passed the trade-off gate when

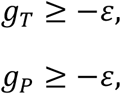

and

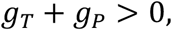

With

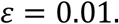

The tolerance allowed one shape metric to worsen by at most 0.01 when the improvement in the other metric was sufficient to produce a positive net gain.

The gate was defined relative to the parent, but candidates passing the gate were ranked according to their absolute shape quality:

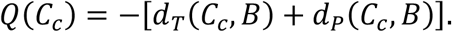

Higher *Q* therefore indicated a state that was globally closer to B, independently of the magnitude of the most recent local step.

Candidates were also required to contain a valid anchor component, contain at least one grown heavy atom connected to the warhead-containing component, have a larger anchor-component heavy-atom count than the parent, and retain every heavy atom of the parent scaffold within 0.5 Å in the child anchor component.

For beam search, structural diversity was enforced among the retained states. Let *G*_*i*_ and *G*_*j*_ denote the coordinates of the grown heavy atoms in two candidates, excluding the seven original warhead atoms. Their symmetric Chamfer distance was

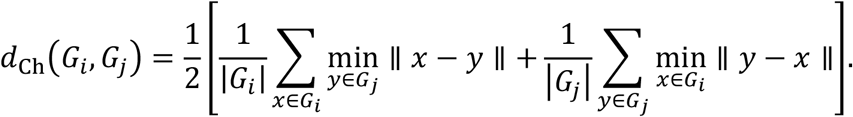

Candidates were considered sufficiently diverse when

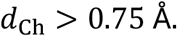

No alignment was performed because all candidates shared the receptor coordinate frame and the fixed warhead.

The target anchor size was defined from the heavy-atom count of ligand B: 16 heavy atoms for the hard pair and 19 for the moderate pair. A candidate graduated when its anchor-component size was within one heavy atom of the target:

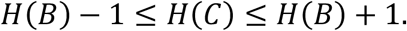

Candidates exceeding *H*(*B*) + 1 were classified as oversize and were not propagated. Smaller candidates remained eligible for further growth.

Two graduation endpoints were recorded, component graduated indicated that the anchor component reached the target size, whereas strictly connected graduated additionally required the complete generated record to form a single heavy-atom component.

A trajectory terminated when every search state had either graduated or died, or when a ceiling of eight stages was reached. The full design comprised two ligand pairs, ten seeds, and two search variants, giving 40 trajectories.

### Adaptive soft-scaffold inpainting and beam search

The adaptive-flexibility campaign fixed the per-stage growth increment at *n*_add_ = 4 heavy atoms and treated the rigidity of the inherited parent scaffold as the search variable. At stage *s*, ligand atoms were partitioned into a hard set *M*_H_ containing the seven-heavy-atom warhead, a soft set *M*_S_ containing atoms retained from preceding stages, and a free set *M*_U_ containing newly generated atoms ^13,14^. At reverse-diffusion step *t*, a stage-specific rigidity coefficient *ρ*_*s*_ ∈ [0,1] controlled how strongly the soft parent coordinates were reintroduced:

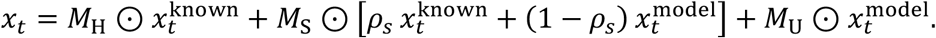

Atom-feature channels remained fixed for all inherited atoms regardless of *ρ*_*s*_, so that the action modulated coordinate flexibility without altering atom types. The warhead remained hard throughout. For *ρ*_*s*_ < 1, the per-sample ligand coordinate mean was subtracted from ligand and pocket coordinates after every reinjection, restoring the zero-centre-of-mass invariant. The *ρ*_*s*_ = 1 branch reproduced the original hard-inpainting path exactly.

Because generated records did not reliably preserve input atom indices, parent identity was recovered by Hungarian assignment constrained by element and coordinate distance. Warhead atoms were required to match within 0.2 Å; soft-parent displacement was unrestricted. Retained children were reordered and inherited atom types were required to remain unchanged.

Each trajectory began from the bare warhead, so stage 1 used only *ρ*_1_ = 1. At every later stage, the action set was

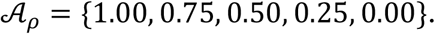

Ten samples were generated per live parent and action, using the same random seed across *ρ* values from a given parent. Sampling used the frozen model with λ = 20, α = 0.3, 50 reverse steps, and five resampling iterations. Greedy search retained one continuation per stage (*k* = 1); beam search retained up to three (*k* = 3). Candidates were filtered, ranked by

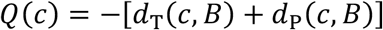

and required to satisfy the same trade-off gate and Chamfer diversity criterion as the variable-increment campaign. The campaign comprised two ligand pairs, ten seeds, and two search variants, yielding 40 independent trajectories.

## RESULTS

### Single-shot shape guidance converts molecular generation into disconnected volume coverage

The seven-heavy-atom warhead remained within the prespecified positional tolerance throughout reverse diffusion in all evaluated conditions. The local inpainting constraint therefore operated as intended, and the structural effects described below did not arise from displacement or loss of the fixed warhead (Fig. 1A-C).

**Figure 1.**
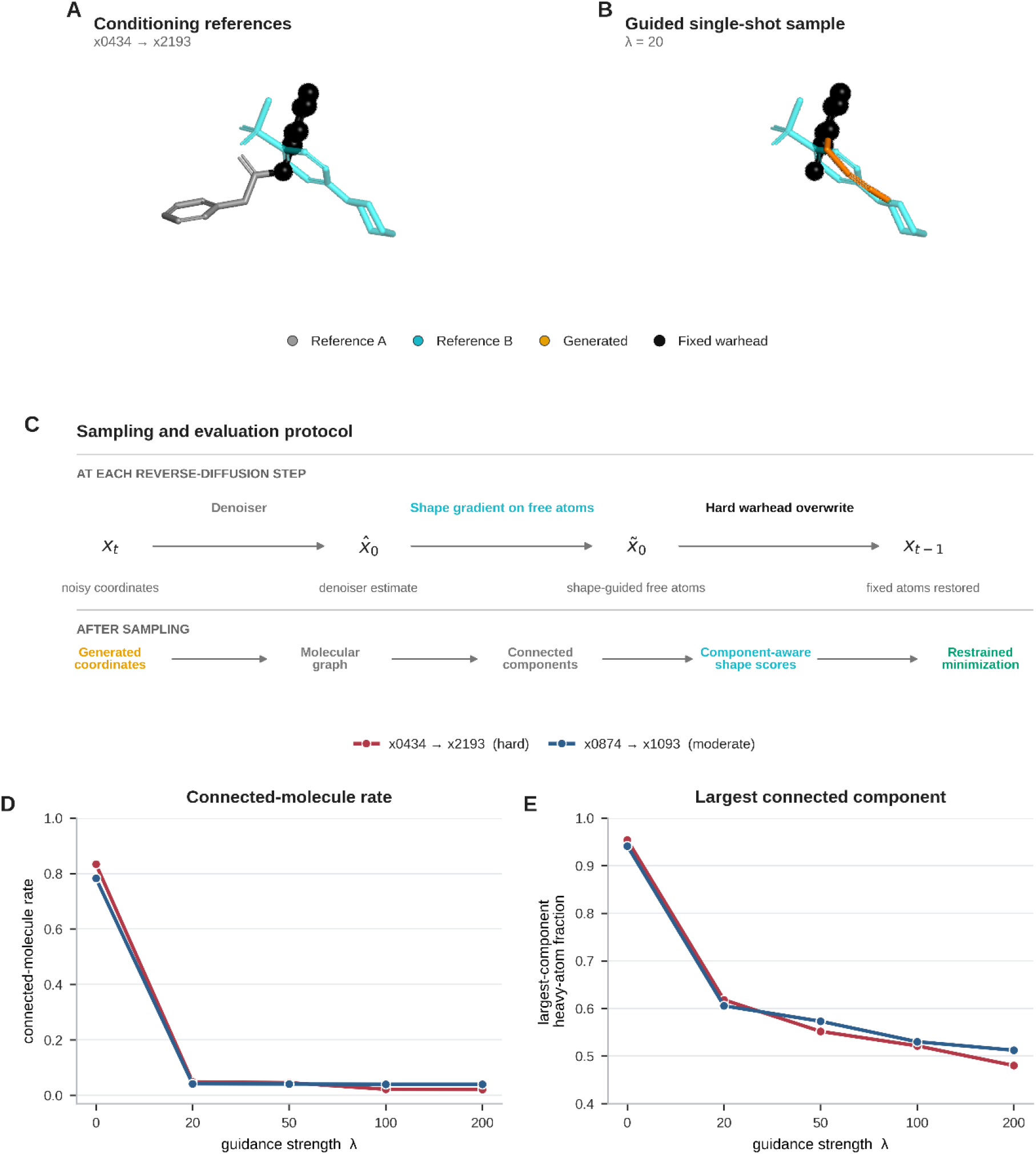
Dual-conditioning task, sampling protocol, and connectivity failure under single-shot shape guidance. **A**, crystallographic conditioning references for the hard x0434-to-x2193 pair in the common receptor coordinate frame. Ligand A provides the fixed seven-heavy-atom warhead, whereas ligand B defines the global shape target. **B**, representative single-shot sample generated with guidance strength λ = 20. **C**, operations performed during each reverse-diffusion step and subsequent structural evaluation. **D**, full-record connected-molecule rate as a function of guidance strength. **E**, mean fraction of generated heavy atoms contained in the largest connected component.

Full-record connectivity responded very differently to the global shape objective. In the absence of guidance, the mean connected-molecule rate was 0.783-0.833 across the two ligand pairs. At the first non-zero guidance strength, λ = 20, it fell to 0.042-0.049 and remained between 0.021 and 0.045 at higher guidance strengths (Fig. 1D). Most of the connectivity loss therefore occurred as soon as global guidance was introduced rather than increasing gradually with λ.

The largest connected component showed the same transition. At λ = 0, it contained 94.1-95.4% of the generated heavy atoms. This fraction decreased to 60.6-61.8% at λ = 20 and to 48.0-51.2% at λ = 200 (Fig. 1E). Because the total atom budget remained unchanged, the reduction reflected redistribution of atoms into secondary disconnected components rather than removal of atoms from the generated record.

Thus, single-shot guidance did not produce a gradual trade-off between connectivity and target-directed remodeling. The first application of the global shape objective was sufficient to convert most generated records from single molecules into collections of disconnected components. This failure motivated reformulating the task as a sequence of smaller scaffold expansions.

### Small staged expansions recover connectivity but do not by themselves impose directional control

To isolate the effect of local growth-action size, we performed a single-step expansion sweep in which every condition began from the bare seven-heavy-atom warhead. Each generated record therefore represented one independent expansion; no resulting scaffold was recursively propagated to a subsequent stage.

Connectivity depended strongly on the number of atoms requested in this single growth action. In the hard pair, the connected fraction decreased from 0.642 when two atoms were requested to 0.098 when eight atoms were requested. The moderate pair showed the same ordering, falling from 0.583 at +2 to approximately 0.03 at +6 and +8. The model-selected auto setting produced connected fractions of 0.050 and 0.124 in the hard and moderate pairs, respectively, and therefore did not recover the connectivity obtained with the smallest fixed increment (Fig. 2A).

**Figure 2.**
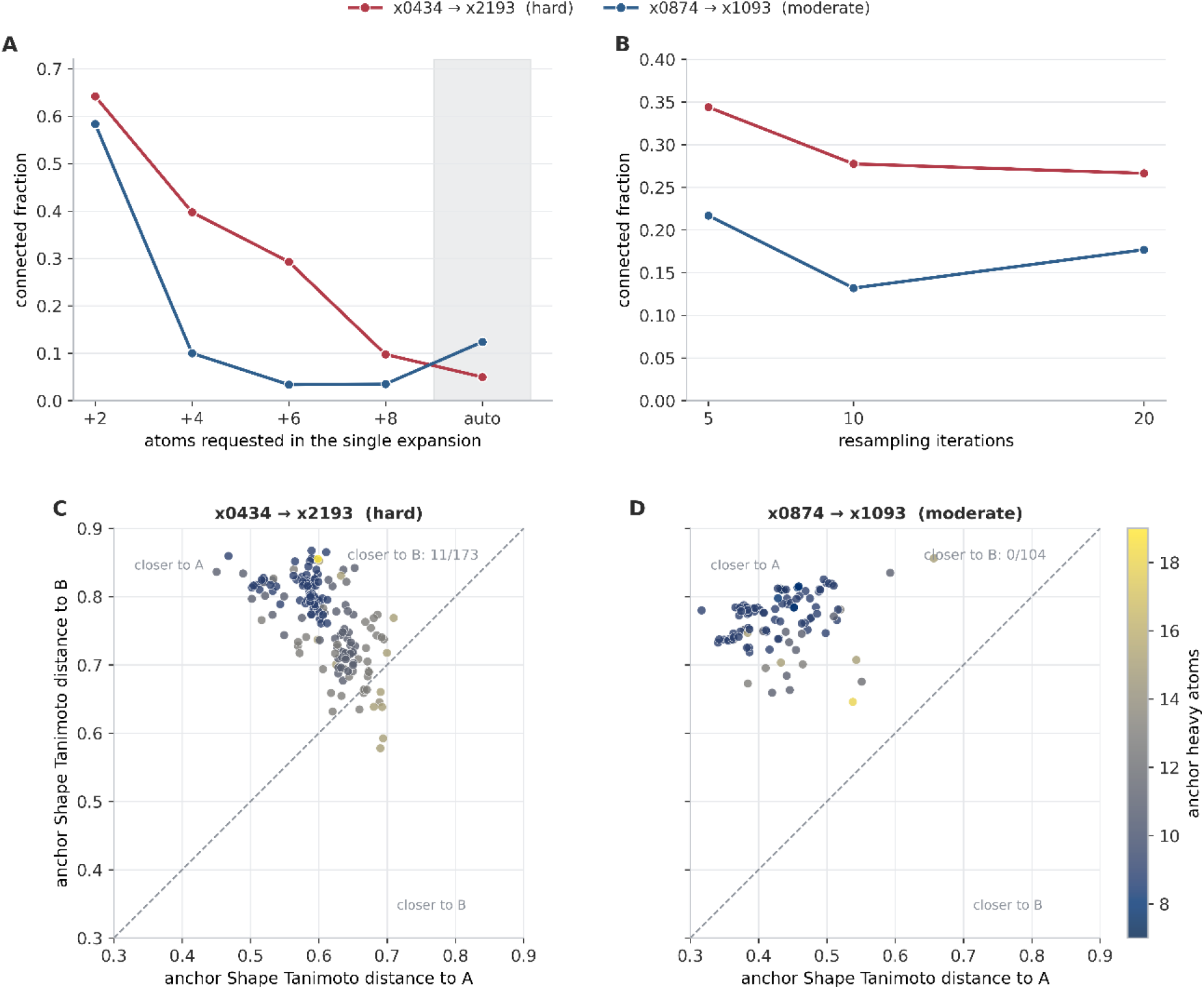
Single-step growth-action size controls connectivity and defines the proposal space for subsequent staged optimization. Every record in this experiment was generated by one independent expansion from the bare seven-heavy-atom warhead; no generated scaffold was propagated to a subsequent stage. **A**, connected fraction as a function of the number of atoms requested in the single expansion. Values are averaged over guidance strengths and resampling settings. The shaded region denotes the model-selected auto condition. **B**, connected fraction as a function of the number of resampling iterations, averaged over increment and guidance conditions. **C** and **D**, connected, shape-auditable anchors from the hard x0434-to-x2193 and moderate x0874-to-x1093 pairs, respectively, represented in the Shape Tanimoto distance-to-A versus distance-to-B plane. Point color denotes anchor-component heavy-atom count. The dashed diagonal indicates equal distance to A and B; points below the diagonal are closer to B. All shape measurements were performed in the common receptor coordinate frame without molecular alignment.

Increasing the number of resampling iterations did not compensate for larger or less stable growth actions. In the hard pair, mean connectivity was 0.344, 0.277, and 0.266 at 5, 10, and 20 resamplings, respectively. The moderate pair similarly showed no monotonic recovery, with connected fractions of 0.217, 0.132, and 0.177 (Fig. 2B). Increasing guidance strength from λ = 10 to λ = 20 further reduced connectivity, from 0.400 to 0.191 in the hard pair and from 0.232 to 0.118 in the moderate pair.

The connected proposals generated by this one-step sweep generally remained closer to A than to B. Only 11 of 173 connected, shape-auditable hard-pair anchors crossed to the B-proximal side of the equal-distance boundary, and none of the 104 moderate-pair anchors did so (Fig. 2C, D). This result does not indicate that shape guidance was inactive. Rather, it shows that a single guided expansion rarely overcame the geometric bias imposed by a warhead originating from A.

The sweep therefore identified a local action-size effect: smaller expansions were substantially more likely to preserve connectivity and generated a viable proposal distribution for subsequent optimization. This observation motivated the next campaign, in which molecular growth was accumulated over multiple stages and repeated sampling and scaffold selection were used to convert favorable proposals into systematic progress toward B.

### B-directed selection outperforms B-blind best-of-ten sampling

The fixed-increment campaign compared three ways of propagating a scaffold through multiple growth stages. In the A1 branch, one candidate was generated at each stage and propagated when it contained a valid anchor component. This branch therefore represented unselected single-sample staged growth. In the A10 branch, ten candidates were generated at each stage, and the valid anchor component containing the greatest number of heavy atoms was propagated without consulting ligand B. In the directed branch, ten candidates were also generated, but propagation explicitly used the shape of B: candidates were required not to worsen either Shape Tanimoto or Shape Protrude distance relative to the parent scaffold, and the candidate providing the greatest joint improvement was selected.

The primary controlled comparison was therefore between A10 and directed selection. Both branches generated ten candidates per stage and differed only in whether ligand B was used to select the continuation scaffold. A1 served as an auxiliary control for the robustness of single-sample sequential growth.

The campaign comprised 560 trajectories. All trajectories yielded an auditable terminal outcome, although some terminated before completing their scheduled number of stages. Thirty-three of the 240 A1 trajectories ended early because their single generated candidate did not provide a valid continuation or because generation failed. None of the 240 A10 trajectories terminated for structural reasons, showing that sampling ten candidates provided a robust fallback when individual proposals were invalid. Three of the 80 directed trajectories ended because none of the ten candidates satisfied the joint shape-progression criterion.

A1 produced the smallest mean improvement toward B at every tested increment in both ligand pairs. Because A1 differed from the other branches in both candidate number and selection opportunity, subsequent analyses focused on the controlled A10-directed comparison.

Directed selection produced greater mean Shape Tanimoto improvement toward B than A10 in all eight ligand-pair-by-increment comparisons (Fig. 3A). The largest difference occurred under the +4 schedule. In the hard pair, mean improvement was 0.195 under directed selection and 0.132 under A10. In the moderate pair, the corresponding values were 0.122 and 0.068.

**Figure 3.**
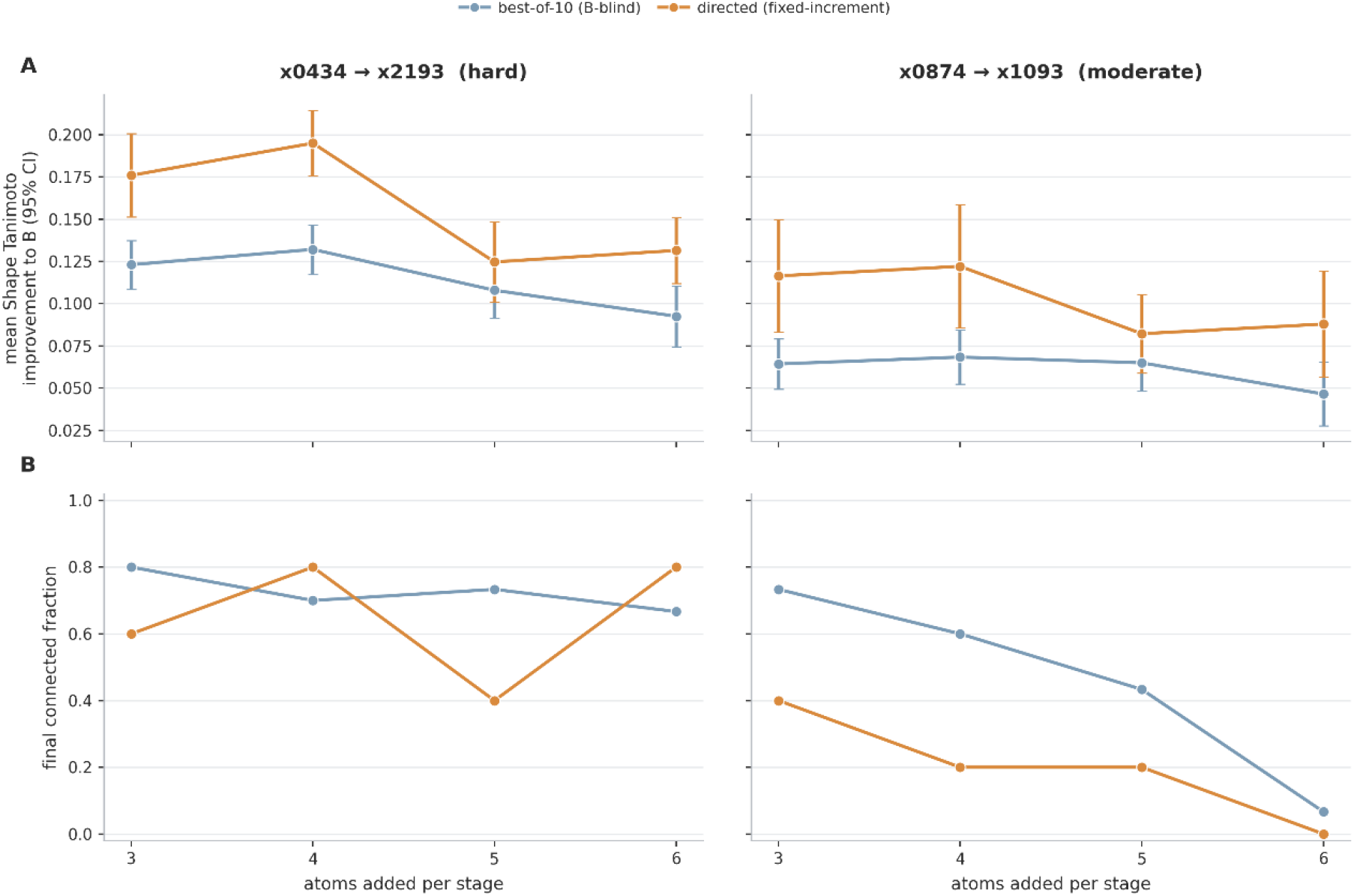
Fixed-increment granularity controls both directional improvement and final connectivity. Columns show the hard x0434-to-x2193 and moderate x0874-to-x1093 pairs. **A**, mean trajectory-level Shape Tanimoto improvement toward B as a function of the fixed number of atoms added per stage. Improvement was calculated as first-stage distance minus final-stage distance, such that positive values indicate movement toward B. Error bars show normal-approximation 95% confidence intervals. **B**, fraction of trajectories whose final generated record was fully connected. The B-blind A10 and directed branches both generated ten candidates per stage and differed only in whether B was consulted during scaffold selection. Each A10 cell contains 30 trajectories and each directed cell contains 10 trajectories.

The directed advantage was statistically supported at +3, +4, and +6 in both ligand pairs under the prespecified one-sided Mann-Whitney test (*p* ≤ 0.015). The difference at +5 was smaller and did not reach the significance threshold in either the hard pair (*p* = 0.178) or the moderate pair (*p* = 0.092).

Among the tested fixed schedules, +4 produced the greatest mean Shape Tanimoto improvement in the directed branch of both pairs. This maximum was pronounced in the hard pair but weaker in the moderate pair, whose response varied less across increments (Fig. 3A). The result identifies +4 as the best-performing schedule within the tested design, rather than establishing a universal optimum for the number of atoms added per stage.

Final connectivity revealed a pair-dependent cost of directional selection. In the hard pair, connectivity varied across increments without a consistent difference between A10 and directed selection. The directed branch reached a connected fraction of 0.80 under both the +4 and +6 schedules, whereas A10 ranged from 0.67 to 0.80 (Fig. 3B).

The moderate pair showed a stronger trade-off. Final connectivity declined with increasing increment size in both branches and was lower under directed selection at every tested schedule. In the directed branch, it decreased from 0.40 at +3 to 0.00 at +6, whereas A10 declined from 0.73 to 0.07 (Fig. 3B). Thus, explicit selection toward B improved shape progression but did not remove the competition between target-directed remodeling and molecular connectivity.

The trajectory-level distributions showed that the directed advantage was not driven by a small number of unusually favorable trajectories. Directed selection shifted Shape Tanimoto improvement upward across every tested increment in both ligand pairs (Fig. 4A). The shift was larger in the hard pair, whereas the moderate pair showed a smaller but consistently positive response.

**Figure 4.**
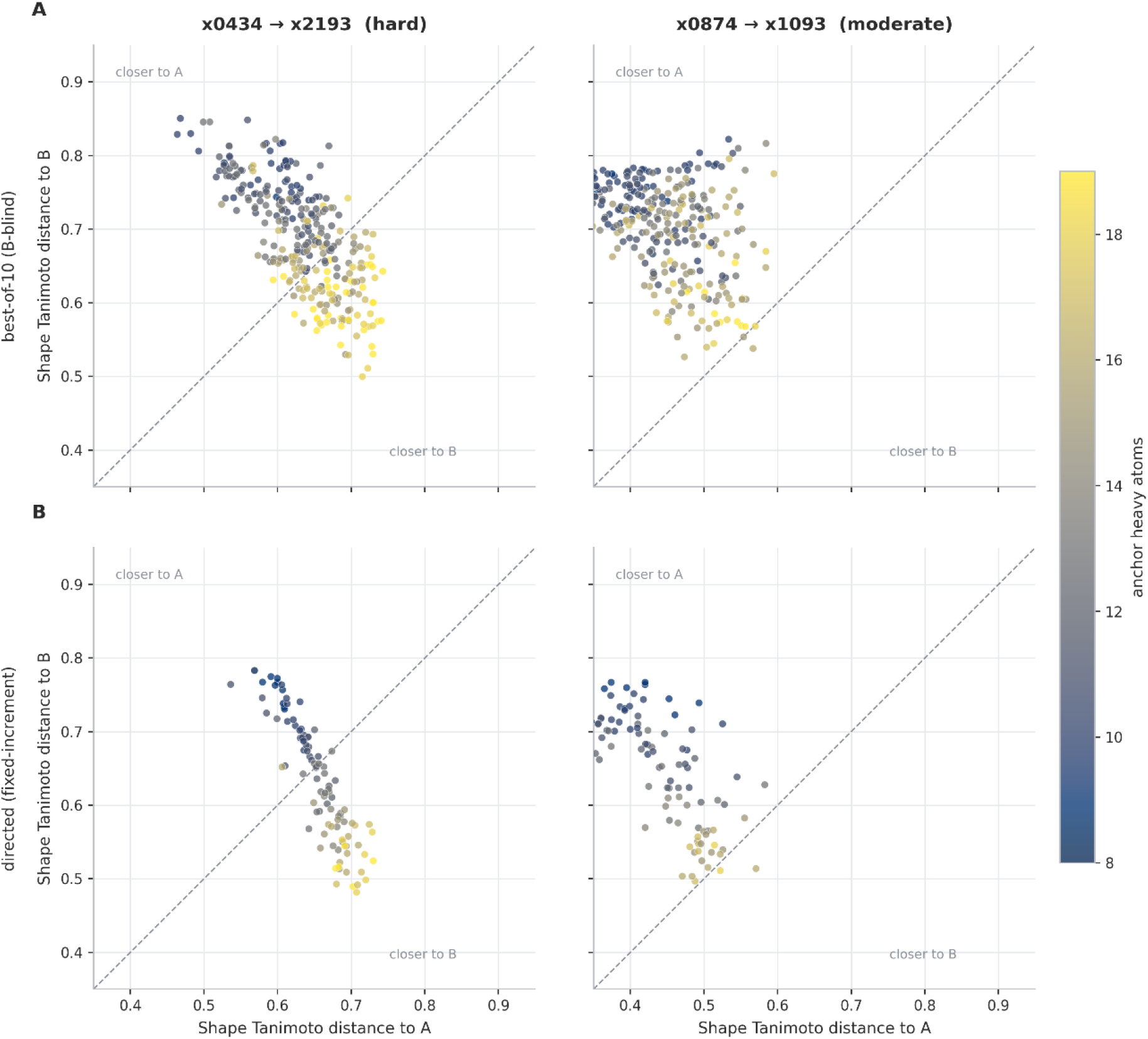
B-directed selection improves global shape overlap and directional target-volume coverage relative to B-blind best-of-ten sampling. **A**, trajectory-level Shape Tanimoto improvement toward ligand B across fixed per-stage increments for the hard x0434-to-x2193 and moderate x0874-to-x1093 pairs. **B**, corresponding improvement in directional Shape Protrude distance to B. Improvement was calculated as the distance at the first generated stage minus the distance at the terminal auditable stage; positive values therefore indicate movement toward B, whereas negative values indicate deterioration.

Shape Protrude revealed a complementary effect that was particularly important in the moderate pair. Mean Protrude improvement under A10 was negative at every increment, ranging from -0.035 to -0.004. Directed selection instead maintained positive mean improvement throughout, ranging from 0.035 to 0.056 (Fig. 4B). Directed selection therefore reduced the fraction of the growing anchor extending outside the target volume even when the change in overall Shape Tanimoto was more modest.

In the hard pair, both A10 and directed selection generally produced positive Protrude improvement, but the directed distributions remained higher at every increment (Fig. 4B). Together, the two metrics show that B-aware selection improved both overall volumetric overlap and directional target-volume coverage, although the relative contribution of each effect differed between ligand pairs.

The A-B distance plane further illustrated these distinct remodeling regimes (Fig. 5). In the hard pair, many fully connected stage states crossed the equal-distance diagonal and became closer to B than to A. Among fully connected hard-pair states, 37 of 48 directed states were B-proximal, compared with 84 of 235 A10 states.

**Figure 5.**
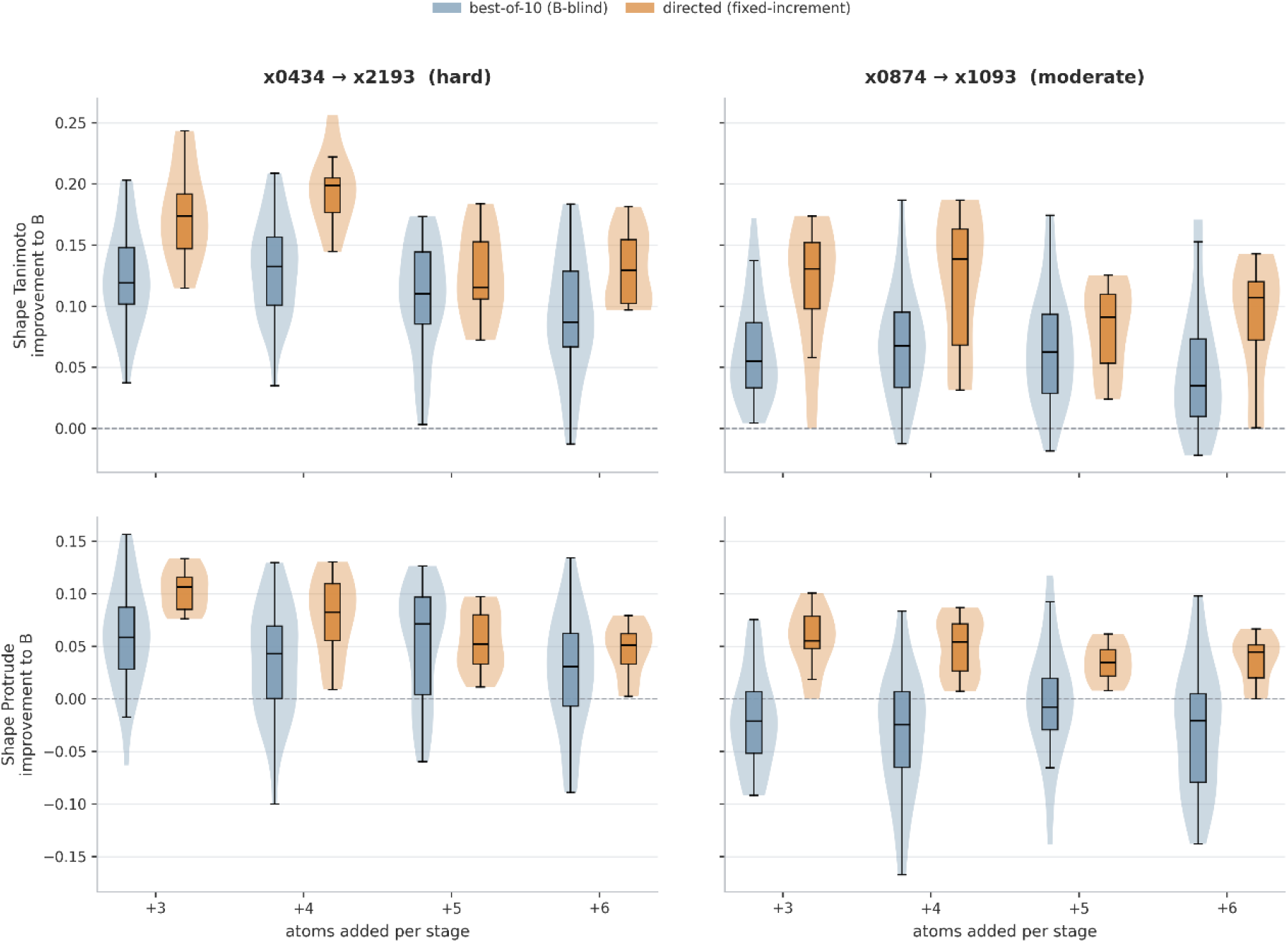
Connected staged-growth states occupy distinct regions of the A-B shape plane under B-blind and B-directed selection. Shape Tanimoto distance to A is plotted against Shape Tanimoto distance to B for fully connected, shape-auditable stage states from the fixed-increment campaign. Columns show the hard x0434-to-x2193 pair and the moderate x0874-to-x1093 pair; rows show best-of-ten B-blind selection (A10) and directed selection. Point color denotes the heavy-atom count of the anchor component. The dashed diagonal indicates equal distance to A and B. States below the diagonal are closer to B, whereas states above the diagonal remain closer to A.

This transition was strongly dependent on scaffold size under directed selection. Among connected directed anchors containing at least 13 heavy atoms, 32 of 36 lay closer to B than to A. The corresponding proportion under A10 was 53%.

Crossing the A-B boundary was rare in the moderate pair. Only 2 of 26 fully connected directed states and 1 of 141 fully connected A10 states were closer to B than to A. Nevertheless, the directed branch shifted moderate-pair states toward B, particularly according to the directional Protrude metric, even though most scaffolds remained more similar to A within the explored size range.

The fixed-increment campaign therefore established three effects within a single experimental design. First, generating multiple candidates per stage greatly increased the robustness of scaffold continuation relative to single-sample growth. Second, explicit B-aware selection produced directional improvement beyond the best-of-ten sampling advantage. Third, the magnitude and structural cost of that improvement depended on both the ligand pair and the fixed growth schedule. These observations motivated treating the number of atoms added at each stage as an adaptive search decision rather than as a fixed parameter.

### Adaptive increment beam search improves endpoint recovery without consistently improving final shape

This campaign was the first direct comparison between greedy search and beam search within the staged-growth framework. Both methods explored the same action space, n_add ∈ {1,2,3,4,5}, and used the same candidate-generation procedure, structural filters, progression gate, and ranking function. The only difference was the number of states retained after each stage. Greedy search propagated a single continuation (k=1), whereas beam search retained up to three structurally diverse alternatives (k=3).

Beam search increased the recovery of strictly connected graduated endpoints. For the hard pair, all 10 seeds produced a strict endpoint under beam search, compared with 6 under greedy search. For the moderate pair, the corresponding numbers were 8 and 3. This suggests that some trajectories selected by greedy search reached dead ends at later stages, while retaining several alternatives allowed the search to continue along a viable route.

The improvement in completion was not accompanied by a consistent improvement in endpoint shape. In the hard pair, the median Shape Tanimoto distance to B decreased from 0.544 under greedy search to 0.523 under beam search. In the moderate pair, the median values were similar, 0.458 for greedy and 0.462 for beam (Fig. 6). The main benefit of the wider beam was therefore greater robustness in reaching a connected target-sized endpoint, rather than systematically producing a better final shape.

**Figure 6.**
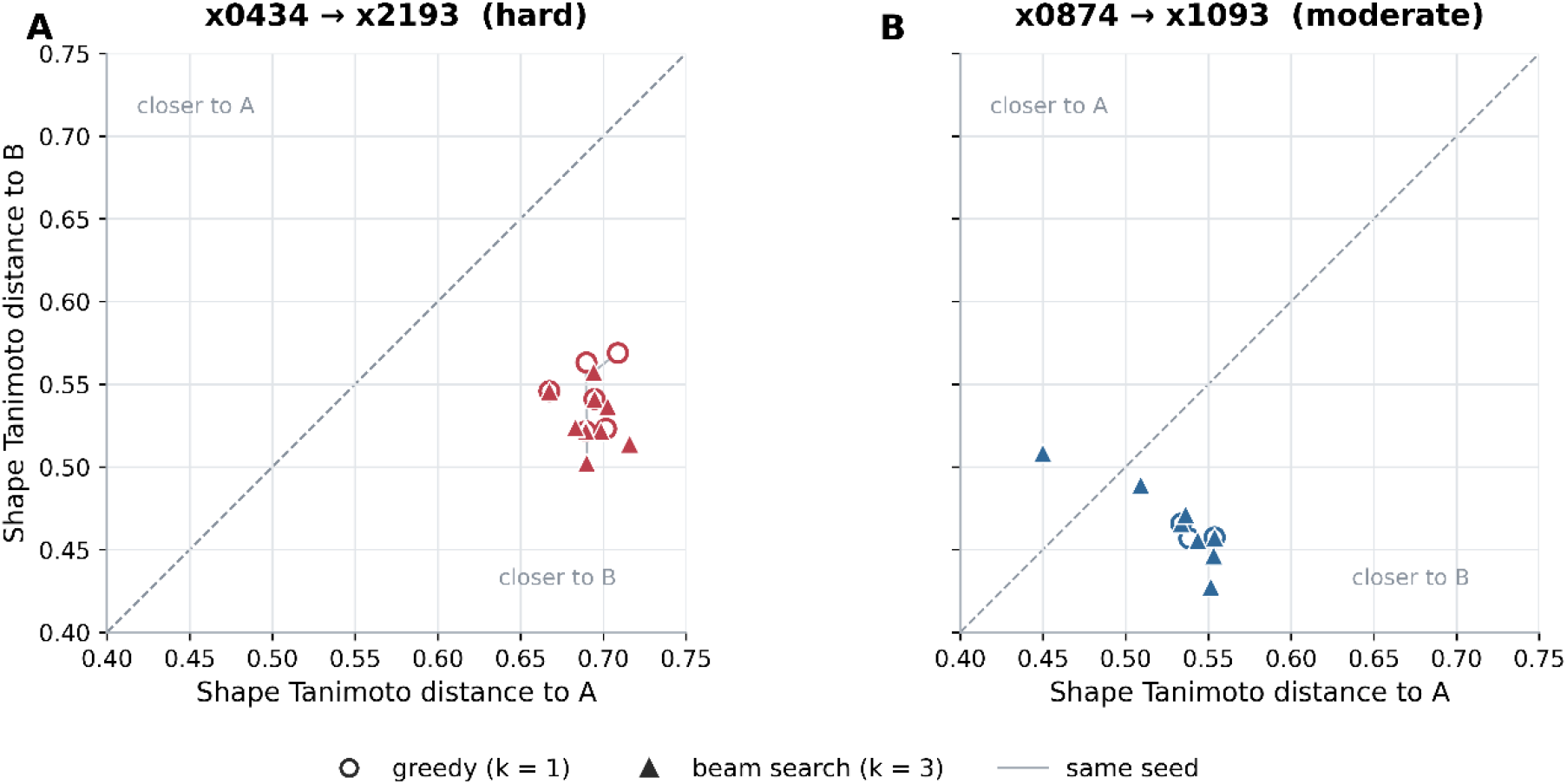
Greedy and beam search under adaptive per-stage increment selection. Shape Tanimoto distance to ligand A is plotted against distance to ligand B for the best strictly connected graduated endpoint obtained from each seed in the hard x0434→x2193 pair (A) and moderate x0874→x1093 pair (B). Both methods selected *n*_add_adaptively from {1,2,3,4,5}and differed only in search width. Open circles denote greedy search (*k* = 1), filled triangles denote beam search (*k* = 3), and grey lines connect endpoints obtained from the same seed. The dashed diagonal indicates equal distance to A and B. Points below the line are more similar to B. Only strictly connected endpoints reaching the target anchor size are shown.

### Adaptive soft-scaffold inpainting and beam search

Adaptive soft-scaffold search produced at least one graduated anchor in 36 of the 40 trajectories, showing that staged growth remained viable when the geometry of the inherited scaffold was allowed to relax. For the hard pair, the median combined Shape Tanimoto and Shape Protrude distance to B was 0.879 under greedy search and 0.866 under beam search, compared with 1.643 for the initial ligand A. For the moderate pair, the corresponding medians were 0.774 and 0.726, compared with 1.228 for A. The adaptive procedure therefore produced target-sized anchors that were substantially closer to B than the starting reference in both systems (Fig. 7A, 7B).

**Figure 7.**
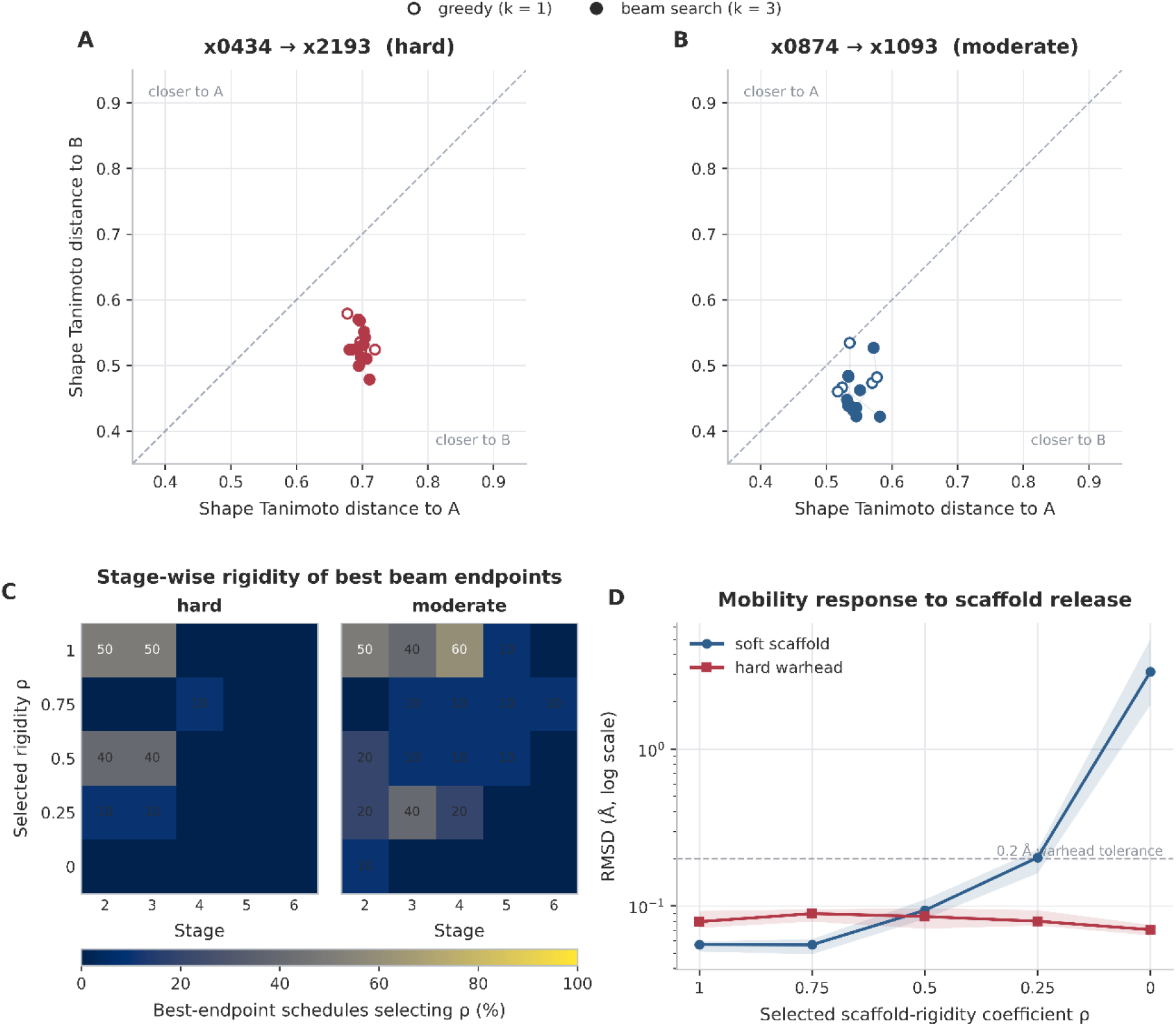
Adaptive control of scaffold rigidity during staged molecular growth. A–B, Best graduated anchor from each successful greedy (*k* = 1, open circles) or beam (*k* = 3, filled circles) trajectory, shown by Shape Tanimoto distance to ligands A and B. Points below the diagonal are more B-like. C, Rigidity coefficient *ρ*selected at each stage along the best beam endpoint from each seed; values show the percentage of trajectories. D, Median soft-scaffold and warhead RMSD versus *ρ*, with interquartile ranges. Lower *ρ*progressively increased scaffold mobility while preserving the fixed warhead below the 0.2 Å threshold.

The rigidity coefficient acted as a graded control of scaffold mobility. Across 194 selected post-initial-stage states, *ρ*was strongly inversely associated with soft-scaffold RMSD (Spearman *ρ* = −0.718). Median displacement increased from approximately 0.02 to 0.05 Å at *ρ* = 1, to 0.05 to 0.07 Å at *ρ* = 0.75, 0.09 to 0.11 Å at *ρ* = 0.5, and 0.15 to 0.22 Å at *ρ* = 0.25. Complete release at *ρ* = 0 was rarely selected and produced displacements of several ångströms (Fig 7D). In contrast, median warhead RMSD remained below approximately 0.11 Å across all groups. The method therefore introduced mobility mainly into the previously generated scaffold while preserving the fixed seven-atom warhead.

Beam search mainly improved the robustness of the adaptive procedure. Completion increased from 9/10 to 10/10 trajectories for the hard pair and from 7/10 to 10/10 for the moderate pair. Final shape quality was similar between methods for the hard pair, with no significant paired difference among the nine common successful seeds (*p* = 0.547). For the moderate pair, beam search produced a lower median combined distance and improved Shape Protrude in all seven paired successful seeds (*p* = 0.0156), although the combined endpoint difference was not significant (*p* = 0.109). Beam search also required one additional stage on average, suggesting that its main contribution was to preserve viable longer trajectories rather than to accelerate convergence.

### Cross-campaign progression toward the target shape

The best observed anchor components showed a progressive reduction in their combined Shape Tanimoto and Shape Protrude distance to ligand B across the successive experimental campaigns (Table 1).

**Table 1.** Best target-directed anchor components obtained across campaigns. *d*_*T*,*B*_and *d*_*P*,*B*_denote Shape Tanimoto and Shape Protrude distances to ligand B, respectively, and *S*_*B*_ = *d*_*T*,*B*_ + *d*_*P*,*B*_. Lower values indicate greater shape similarity to B; reduction from A was calculated from *S*_*B*_.

| Pair | Method | Schedule | $d_{T,B}$ | $d_{P,B}$ | $S_B$ |
| --- | --- | --- | --- | --- | --- |
| Hard | Initial A | — | 0.874 | 0.769 | — |
| Hard | Single-step +4 | +4 | 0.678 | 0.406 | 34.0% |
| Hard | Directed fixed +4 | +4→+4→+4 | 0.493 | 0.334 | 49.7% |
| Hard | Adaptive growth | variable | 0.487 | 0.320 | 50.9% |
| Hard | Adaptive mask | variable,<br>$n_{\text{add}} = 4$, | 0.479 | 0.337 | 50.4% |
| Moderate | Initial A | — | 0.728 | 0.500 | — |
| Moderate | Single-step +4 | +4 | 0.659 | 0.380 | 15.4% |
| Moderate | Directed fixed +4 | +4→+4→+4 | 0.503 | 0.311 | 33.7% |
| Moderate | Adaptive growth | variable | 0.427 | 0.283 | 42.1% |
| Moderate | Adaptive mask | variable,<br>$n_{\text{add}} = 4$, | 0.422 | 0.275 | 43.2% |

For the hard x0434→x2193 pair, the combined Shape Tanimoto and Shape Protrude distance decreased from 1.643 for ligand A to 1.084 after the best single-step +4 expansion and to 0.827 for the best B-directed fixed-+4 trajectory. Allowing *n*_add_to vary adaptively produced a further reduction to 0.806, corresponding to a 50.9% improvement relative to A and representing the best combined result for this pair. Adaptive soft-scaffold search reached a similar value of 0.816, with a slightly lower Shape Tanimoto distance than adaptive growth, 0.479 versus 0.487, but a higher Shape Protrude distance, 0.337 versus 0.320 (Fig. 8A). Thus, most of the improvement arose from accumulated B-directed growth, while adaptive control of growth size provided a modest additional advantage.

**Figure 8.**
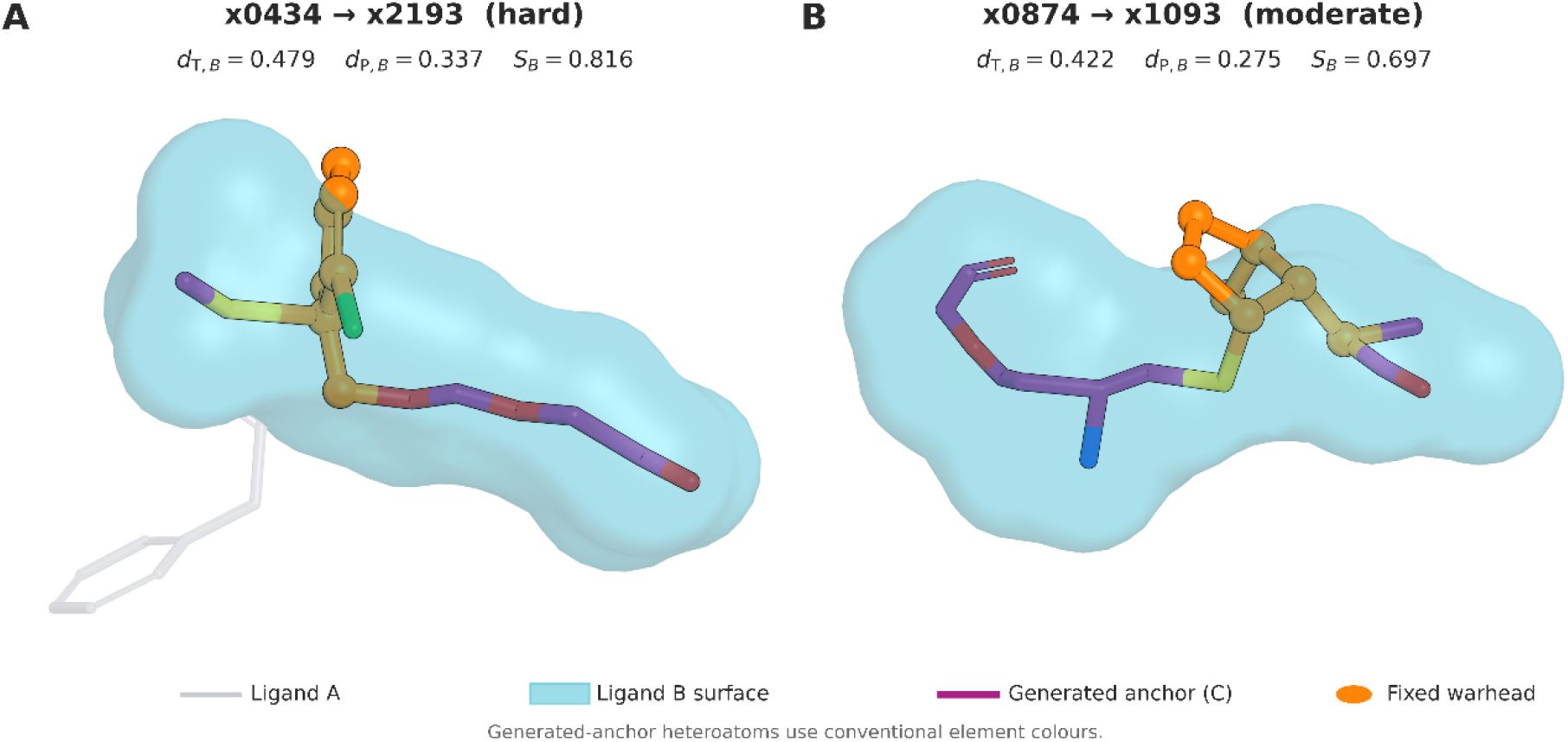
Best target-directed structures obtained by adaptive soft-scaffold beam search. **A**, Best graduated anchor for the hard x0434→x2193 pair. **B**, Best graduated anchor for the moderate x0874→x1093 pair. Ligand A is shown as semitransparent grey sticks, ligand B as a cyan molecular surface, and the generated anchor as sticks with magenta carbon atoms and conventionally coloured heteroatoms. The fixed seven-heavy-atom warhead is highlighted in orange. Structures are displayed in the common receptor coordinate frame without post hoc alignment. Reported values correspond to the Shape Tanimoto distance to B (*d*_*T*,*B*_), Shape Protrude distance to B (*d*_P,*B*_), and their sum (*S*_*B*_).

For the moderate x0874→x1093 pair, the combined distance decreased from 1.228 for ligand A to 1.039 after the best single-step +4expansion and to 0.814 under directed fixed-+4 growth. Adaptive *n*_add_search reduced this value to 0.711, with Shape Tanimoto and Shape Protrude distances of 0.427 and 0.283, respectively. Adaptive soft-scaffold search produced the best endpoint for this pair, further reducing the combined distance to 0.697 and the individual distances to 0.422 and 0.275. These results correspond to reductions of 42.1% and 43.2% relative to A for adaptive growth and adaptive masking, respectively.

The comparison suggests that adaptive growth granularity benefited both systems, whereas controlled scaffold mobility provided an additional advantage mainly for the moderate pair. Cross-campaign comparisons remain descriptive because sampling budgets, trajectory lengths, and search spaces differed between experiments.

## DISCUSSION

The utility of diffusion models rests on their capacity to construct ligands as a function of their geometric pocket, given the training distribution and the learned prior. In the present work we applied a geometric force to the denoising process of a frozen DiffSBDD model in order to grow a warhead derived from a molecule A sequentially toward the shape of a molecule B, through discrete growth stages. In this way, the diffusion model retained access to the complete molecular context from the outset while sequentially adapting toward the optimal generation.

Staged growth was introduced because of the inability of DiffSBDD to generate plausible, connected molecules at the guidance strength required to steer generation. Full-record connectivity was already lost at λ = 20, where the connected-molecule rate fell from 0.783–0.833 in the absence of guidance to 0.042–0.049, and the fraction of heavy atoms contained in the largest connected component decreased monotonically up to λ = 200 (Fig. 1D, E). A plausible explanation for this failure is that the frozen model was trained to populate a pocket in a single denoising pass under a fixed atom budget, rather than to displace an already seeded atom cloud toward an external target shape. When strong guidance was applied from the warhead, satisfying the volumetric objective by placing loose atoms across the region occupied by B was less costly for the sampler than reorganizing a connected molecular graph, because the shape objective acted on coordinates while connectivity was only inferred post hoc from interatomic distances. Nothing in the sampling trajectory penalized disconnection. This interpretation is consistent with the observation that the effect depended on the geometry of the initial warhead relative to the final shape of B, and that connectivity improved when the number of atoms requested per generative action was reduced rather than imposed as a single final count (Fig. 2A). Staged growth resolved this failure because each expansion was small and local relative to an already connected scaffold, returning the model to the regime for which it was trained, namely filling a limited volume around a fixed structure, so that guidance had to bias only a few atoms at a time. This observation motivated the development of the staged fragment-growth strategy.

This mode of operation differs from that of AutoFragDiff, which performs autoregressive fragment-based elaboration without knowledge of the final target shape and requires explicit training on retrosynthetic growth. Our approach instead conditioned generation on that final shape from the outset and obtained staged growth without retraining, by applying inference-time guidance to a frozen general-purpose generator. The trade-off is that reusing a pretrained model avoids any dedicated training cost but pushes the sampler away from the region of the prior where its outputs are most reliable, which is precisely the failure that staged growth was designed to contain. The soft-masking mechanism additionally allowed a controlled degree of scaffold flexibility that autoregressive elaboration does not provide, and that could in future be exploited to refine scaffolds over iterative cycles, opening a path toward a genuinely hybrid growth mechanism.

In the first staged-growth experiment we observed that, although the optimal range was governed by the geometry of each pair, the best-performing schedule lay near four atoms per stage (Fig. 3A). This effect was amplified when we additionally implemented directed selection (Figs. 3–5), in which ten candidates were generated at each stage and two propagation rules were contrasted: a directed branch that propagated the candidate most similar to B, and a B-blind branch that propagated the anchor component with the greatest number of connected heavy atoms. Directed selection produced greater mean Shape Tanimoto improvement toward B than B-blind best-of-ten sampling in all eight ligand-pair-by-increment comparisons, with the largest difference under the +4 schedule (0.195 versus 0.132 in the hard pair; 0.122 versus 0.068 in the moderate pair; Fig. 3A). The Tanimoto and Protrude criteria were therefore able to outperform selection of the largest connected component across the different levels of granularity (Fig. 4A, B).

The magnitude of this improvement, and the cost it imposed on connectivity, depended on the ligand pair. Directed growth accounted for most of the target-shape progression in the hard pair, whereas controlled scaffold mobility provided an additional advantage in the moderate pair. This pair dependence should be interpreted with caution. Shape Tanimoto and Shape Protrude quantify volumetric disagreement and directional protrusion, but they may not fully capture which shapes of B are geometrically accessible from a given warhead, and therefore may be imperfect predictors of pair difficulty. A more thorough characterization of what makes a target shape reachable, beyond these two distances, would help clarify when directed staged growth is expected to succeed.

Finally, greedy search and beam search were applied to evaluate the roles of granularity and flexibility during generation. The per-stage increment was swept from one to five atoms as an adaptive search decision, and separately, different fractions of the diffusion steps were released on the inherited scaffold to assess whether the structural rigidity of earlier stages was constraining the guidance. The intent was not to establish a difference between the two search strategies but to show that search-based optimization is useful in general within this framework. Both approaches yielded molecules more similar to B, and beam search increased the recovery of strictly connected graduated endpoints without consistently improving final shape.

The natural next step is to move from these search-based optimization procedures toward reinforcement learning architectures, in which the growth policy is learned rather than enumerated stage by stage. A systematic evaluation of the hard hyperparameters that shape the present procedure would also be valuable, in particular the Gaussian width α, which sets the scale of the overlap field, together with the gradient clipping threshold and the Chamfer diversity criterion, each of which currently constrains the behavior of the search in ways that were fixed rather than optimized.

## CONCLUSION

Geometric guidance applied to a frozen pocket-conditioned diffusion model steered generation toward a target shape but did so by producing disconnected molecules, converting single-shot generation into volumetric coverage rather than connected molecular growth. Reformulating the task as a sequence of small scaffold expansions, together with control of the per-stage growth granularity, substantially mitigated this fragmentation, although it did not eliminate the competition between target-directed remodeling and connectivity. The model was able to generate molecules conditioned toward a shape B while efficiently maintaining the inpainting of a warhead A, and it did so most effectively when greedy and beam search were used to optimize growth as a function of fragment size and of growth flexibility, the latter controlled through soft-masking by releasing different numbers of diffusion steps. The success of generation appeared to be closely tied to the geometry of the pairs, which points to a bottleneck in their selection that lies beyond the behavior of the model itself. A logical direction would be to transition from search-based optimization toward reinforcement learning architectures in order to navigate the vast growth landscape, which appears to be governed by the size of the growth increment and by flexibility, and which would likely require a deeper understanding of the roles of α, gradient clipping, and the Chamfer criterion.

## DATA AND CODE AVAILABILITY

The source code, DiffSBDD inference patches, campaign launchers, tests, and reproducibility instructions are available at https://github.com/Ismaelcasku/dual-conditioning under release v2.0.0 and are archived on Zenodo at https://doi.org/10.5281/zenodo.21458637.

The complete generated structures, trajectory-level records, campaign manifests, run-status files, and validation outputs are available in the Zenodo dataset *Dual-conditioning campaigns for pretrained 3D molecular diffusion*, version 2.0.0, at https://doi.org/10.5281/zenodo.21468756. The pretrained DiffSBDD checkpoint and externally sourced structural inputs are not redistributed and must be obtained from their original sources as described in the repository documentation.

## AUTHOR CONTRIBUTIONS

I.C.: Conceptualization, Methodology, Software, Validation, Formal analysis, Investigation, Data curation, Visualization, Writing – original draft, Writing - review and editing.

